# Deep Learning Model Imputes Missing Stains in Multiplex Images

**DOI:** 10.1101/2023.11.21.568088

**Authors:** Muhammad Shaban, Wiem Lassoued, Kenneth Canubas, Shania Bailey, Yanling Liu, Clint Allen, Julius Strauss, James L Gulley, Sizun Jiang, Faisal Mahmood, George Zaki, Houssein A Sater

## Abstract

Multiplex staining enables simultaneous detection of multiple protein markers within a tissue sample. However, the increased marker count increased the likelihood of staining and imaging failure, leading to higher resource usage in multiplex staining and imaging. We address this by proposing a deep learning-based <u>MA</u>rker imputation model for multipleX <u>IM</u>ages (MAXIM) that accurately impute protein markers by leveraging latent biological relationships between markers. The model’s imputation ability is extensively evaluated at pixel and cell levels across various cancer types. Additionally, we present a comparison between imputed and actual marker images within the context of a downstream cell classification task. The MAXIM model’s interpretability is enhanced by gaining insights into the contribution of individual markers in the imputation process. In practice, MAXIM can reduce the cost and time of multiplex staining and image acquisition by accurately imputing protein markers affected by staining issues.

## Introduction

Multiplex staining and imaging, a state-of-the-art technology, has revolutionized the simultaneous visualization of multiple protein markers within a single tissue sample. Various techniques have emerged to capture multiplex images with up to 100 markers, enabling a deeper understanding of complex biological processes (1–5). However, the limited availability of human tissues and the increased number of markers introduce novel challenges, such as a higher likelihood of staining failure (missing or aberrant stain), leading to increased costs and time required for multiplex image acquisition. Additionally, the augmented marker count also amplifies the probability of encountering markers with latent biological relationships. We hypothesized that these relationships could be leveraged to impute protein markers with missing stains in multiplex images, ultimately reducing the necessity for tissue restaining and saving associated extra cost and time.

Artificial intelligence (AI) has demonstrated remarkable success across diverse fields, including medical image analysis (6–10). The field of multiplex imaging has also harnessed the power of AI for tasks such as cell segmentation (11, 12), cell classification (13–15), and spatial analysis (16, 17). The imputation of missing protein markers is a missing data imputation problem, and a number of AI-based methods, particularly deep learning methods, have been developed for missing data imputation in various domains (18–22). However, current literature on image imputation in medical imaging primarily focuses on radiology datasets (20–22), with limited research exploring the potential of deep learning models for marker synthesis in multiplex images (23, 24).

We present a deep learning-based <u>MA</u>rker imputation model for multipl<u>eX IM</u>ages (MAXIM) that harnesses the capabilities of deep learning to accurately impute a protein marker by leveraging the latent relationships between markers in multiplex images (**Figure 1**). We extensively evaluate the MAXIM’s performance at both the pixel and cell levels in whole slide multiplex images as well as specific regions of interest, providing a comprehensive understanding of its imputation capabilities across various cancer types. Additionally, we examine the MAXIM’s performance in a downstream task of cell classification to demonstrate its practical utility beyond imputation. Lastly, we enhance the interpretability of the MAXIM model by investigating the attribution of input markers through aggregated gradients (25), unveiling valuable insights into the contribution of each input marker in the imputation process.

**Fig. 1.**
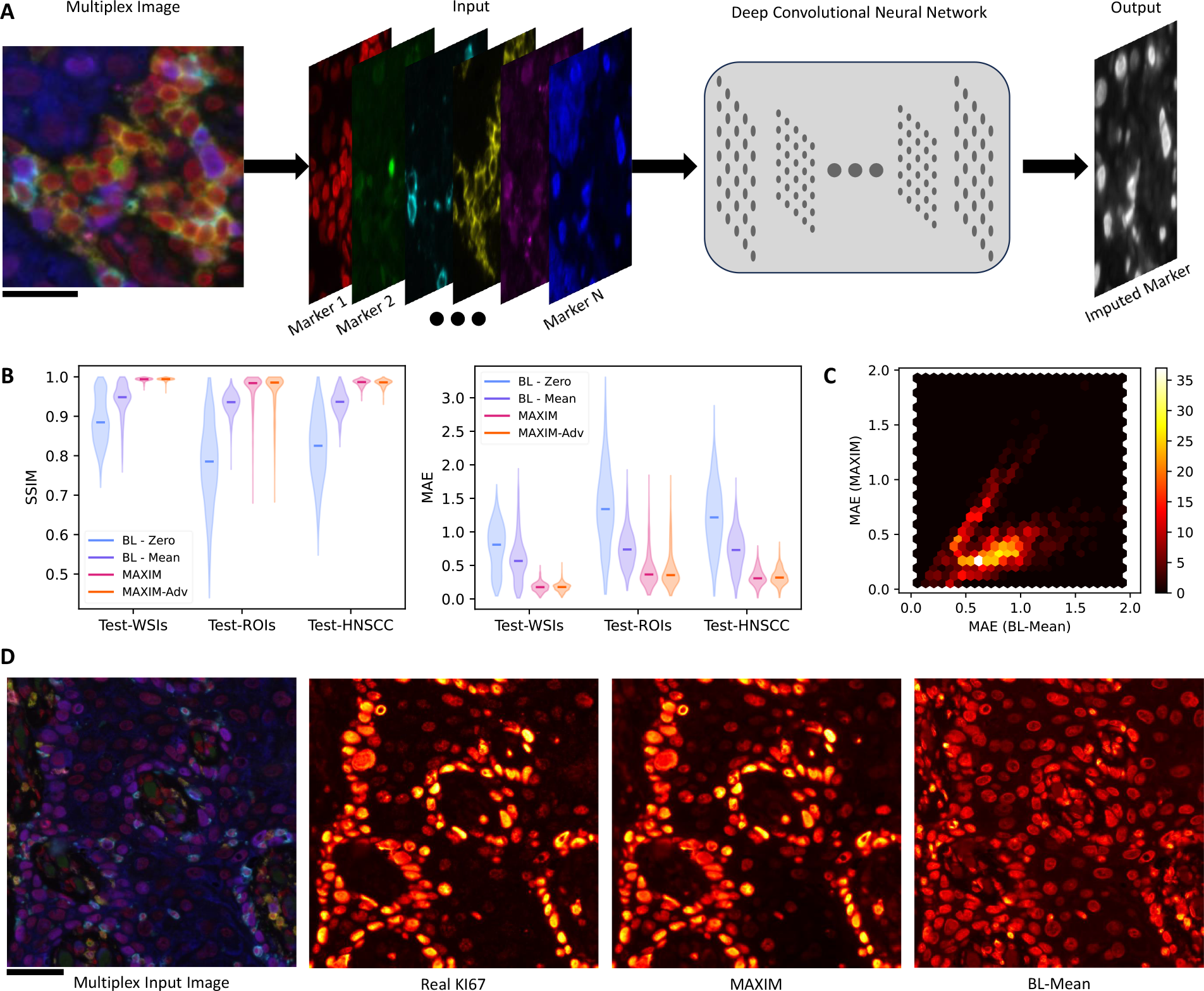
The flow diagram of the proposed method, along with quantitative and visual results for KI67 marker imputation. **(A)** The proposed model (MAXIM) takes a multiplex image with *N* markers as input to impute a marker of interest. Scale bar: 20 *µm*. **(B)** The quantitative assessment of MAXIM imputation using the structural similarity index (SSIM) and mean absolute error (MAE) metrics across three test sets. The width of each violin represents the density of data points (images) with corresponding SSIM/MAE scores, and the solid line within each violin indicates the median SSIM/MAE score. BL-Zero and BL-Mean are pseudo models used to provide baseline MEA and SSIM scores in multiplex images with high sparsity and structural similarity across markers. BL-Zero generates an imputed image with zero values, while BL-Mean creates an imputed image using the mean values of the input markers. MAXIM-Adv is a variant of MAXIM trained with adversarial loss. **(C)** The hexbin plot shows the relationship between the MAE scores of BL-Mean and MAXIM models on the Test-ROIs set. Color intensity reflects the density of data points within each hexagon. **(D)** Visual results of KI67 marker imputation using BL-Mean and MAXIM models for an examplar input image from Test-HNSCC set. The input image consists of six markers: DAPI (red), FOXP3 (green), CD4 (yellow), CD8 (cyan), PDL1 (magenta), and CK (blue). Scale bar: 50 *µ*_*m*_.

### Results

MAXIM was trained and evaluated on a whole slide multiplex immunofluorescence (MxIF) imaging dataset, encompassing cases from four different cancer types: Urothelial, Anal, Cervical, Head and Neck Squamous Cell Carcinoma (HNSCC). A separate MAXIM model was trained for each marker in MxIF images, using the remaining markers as input. The model’s performance is evaluated on images of size 1396 *×* 1860 pixels from three distinct test sets: Test-WSIs (1920 images from 4 HNSCC whole slide multiplex images), Test-ROIs (1097 images from 9 cases of different cancer types), and Test-HNSCC (623 images from 13 cases of HNSCC).

MAXIM performance is evaluated using structural similarity index (SSIM) and mean absolute error (MAE) between the imputed marker images and corresponding real marker images. MAXIM achieved high median SSIM and low median MAE scores (**Figure 1B** and **Extended Data Figure 1**). The results of two baseline pseudo models (BL-Zero, and BL-Mean) were included to contextualize MEA and SSIM scores in multiplex images with high inherent sparsity and structural similarity across markers. The BL-Zero model utilized zero-valued images as imputed images, while the BL-Mean model employed mean images of the input markers as imputed images. MAXIM and its variant MAXIM-Adv, trained with an adversarial loss, exhibited higher performance over baseline results for KI67 marker imputation (**Figure 1B**), particularly on Test-WSIs and Test-HNSCC sets. However, BL-Zeros and BL-Mean also exhibit higher SSIM scores and lower MAE scores in certain images, particularly those with limited tissue structure leading to high sparsity. To conduct a more detailed performance analysis between BL-Mean and the MAXIM models in these particular cases, we employed hexbin plots to compare the paired mean absolute error (MAE) for KI67 marker imputation on the Test-ROI set. The results show that the MAXIM model’s MAE scores are either lower or comparable to those of the BL-Mean model (**Figure 1C**). **Figure 1D** and **Extended Data Figure 2** visually exemplify the MAXIM’s ability to accurately impute the markers of interest. These results highlight the effectiveness of MAXIM in leveraging latent correlations and patterns to accurately impute a marker of interest.

Next, we proceed to evaluate the performance of MAXIM at the cell level on subsets of the Test-ROIs and Test-HNSCC sets (**Figure 2A & Extended Data Figure 3**). These subsets comprise a substantial number of cells, totaling more than 400,000, extracted from 245 images. To assess MAXIM’s performance accurately, we compare the mean cell expression of each cell in real images with corresponding cells in the imputed images. Again, MAXIM shows lower MAE compared to the two baseline methods on both Test-ROIs and Test-HNSCC subsets. Notably, the results obtained at the cell level align closely with the performance patterns observed at the pixel level when evaluated on the same set of images (**Figure 2B & Extended Data Figure 3**). Furthermore, we evaluate the utility of imputed images in a downstream task of cell classification. To establish ground truth cell labels, we utilized the HALO software to categorize cells as either positive or negative based on the mean cell expression of the KI67 marker in real images. Subsequently, we generated a precision-recall curve and determined the average precision by comparing the mean cell expression of each cell in the imputed images with the corresponding cell label in the real images (**Figure 2C**). MAXIM exhibited high average precision in both test sets, affirming its capability in precisely imputing marker values that can be reliably used for downstream analysis.

**Fig. 2.**
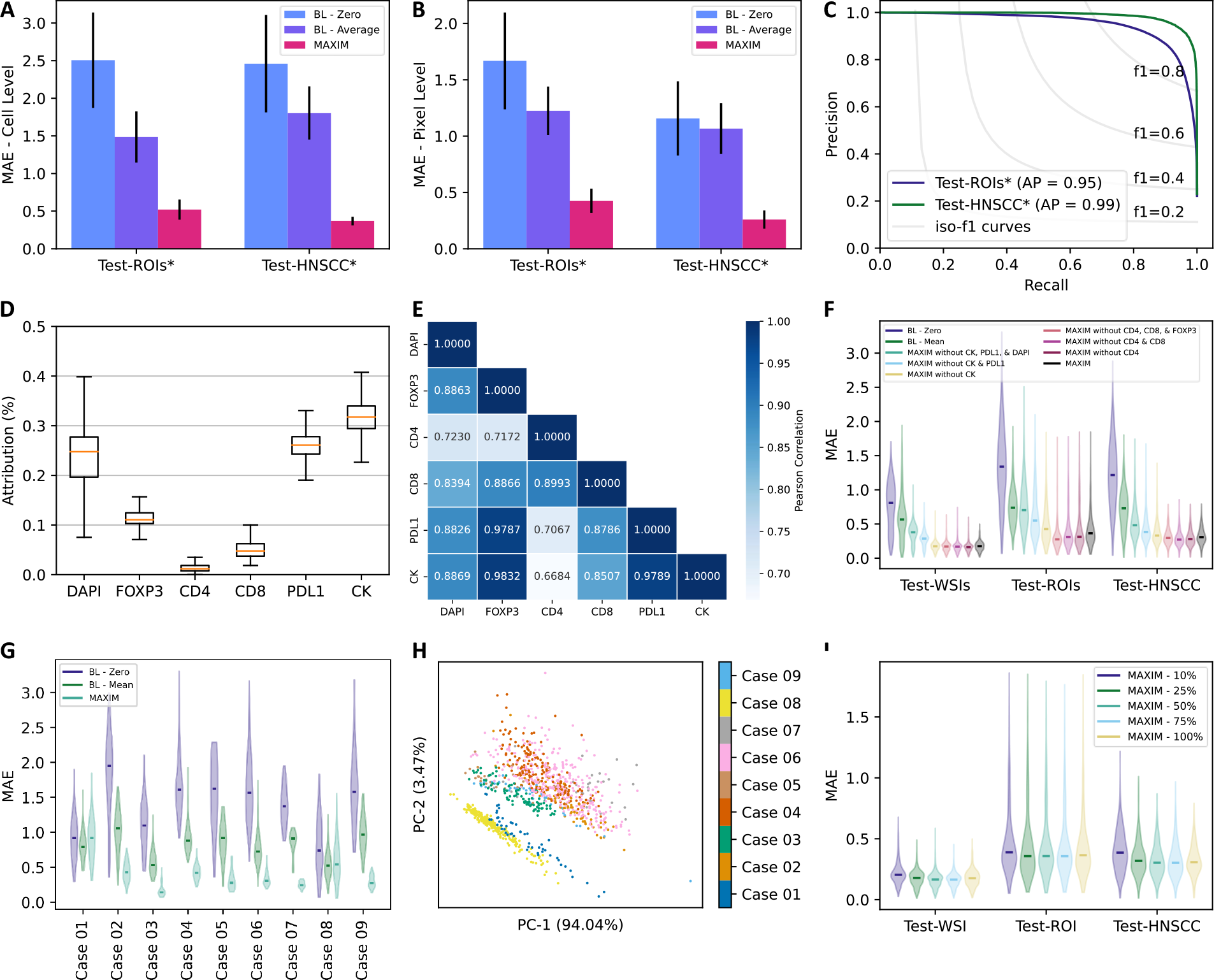
Performace and interpretability analysis of MAXIM. **(A-B)** Evaluation of MAXIM at cell and pixel level on subsets of Test-ROIs and Test-HNSCC sets. The difference in mean expression of each cell is used for cell level evaluation. The height of the bars indicates the average absolute error, whereas error bars represent the standard deviation. **(C)** The precision-recall curve for classifying KI67 cells as positive or negative in MAXIM-based imputed images. The ground truth labels for positive and negative cells were determined using real KI67 marker images. **(D)** The boxplots show the distribution of gradient-based attribution/contribution of each marker towards the KI67 marker imputation. The line inside the box represents the median, the edges of the box represent the interquartile range, and the whiskers extend to 1.5 times the interquartile range to represent the range of the attribution values. **(E)** Pearson correlation between attribution values of input marker pairs. **(F)** Comparison of MAXIM model’s performance using violin plots when trained only using markers with high attribution and low attribution for KI67 marker imputation. The width of each violin represents the density of data points (images), and the solid line within each violin indicates the median attribution. **(G)** Evaluation of MAXIM at case level on Test-ROIs set. **(H)** Two-dimentional projection of all input images for KI67 marker inputation in Test-ROIs set. The 2D PCA projections are calculated from the mean intensities of each marker in the input images. **(I)** Evaluation of MAXIM models when trained using different subsets of the training set.

To investigate MAXIM’s ability to leverage latent relationships among markers for accurate imputation, we assess the attributions of input markers for marker imputation using the aggregated gradient (25) method on the Test-WSIs set (**Figure 2D & Extended Data Figure 4**). The analysis reveals that CK, PDL1, and DAPI markers exhibit higher contributions to KI67 marker imputation than CD8, CD4, and FOXP3 markers. Additionally, the high correlation in attribution scores between pairs of input markers suggests consistent attribution patterns across images (**Figure 2E**). We train separate models excluding markers with high attribution scores for KI67 marker imputation to further validate these attributions. The results in **Figure 2F** demonstrate a performance decrease in models trained without markers with high attribution scores, while models without markers exhibiting low attribution scores consistently perform well. These findings indicate that input markers with high attribution scores possess latent relationships effectively exploited by MAXIM for accurate KI67 marker imputation.

The investigation of MAXIM’s performance at the case level reveals the potential reason for the relatively high variance in its performance on the Test-ROIs set. MAXIM fails to accurately impute markers in two cases, as shown in **Figure 2F & Extended Data Figure 5**. To further explore the reasons behind these failures, we represent each image by calculating the mean expression of each input marker. By using the mean expression of all images for principal component analysis (PCA) and plotting the top two components, we visualize the differences in the distribution of cases (**Figure 2G & Extended Data Figure 6**). Interestingly, the two cases where MAXIM fails to perform well appear as a separate cluster, especially for the KI67 marker, indicating that these cases exhibit distinct distributions compared to the remaining cases in the Test-ROIs set. Finally, we demonstrate that MAXIM can maintain its high performance even when trained using a limited dataset, as illustrated in **Figure 2H**. This finding highlights the robustness of MAXIM, indicating that it can still effectively impute marker values even with a smaller training dataset.

## Discussion

We demonstrate the effectiveness of MAXIM, a deep generative model, in marker imputation for multiplex images. Specifically, we show that MAXIM can accurately impute the values of a protein marker with pixel-level precision by leveraging the latent relationships between available markers. Additionally, we showcase the utility of imputed marker images for cell classification, a common downstream task in multiplex image analysis. Furthermore, we present the analysis of individual marker importance in MAXIM marker imputation process.

The effectiveness of the MAXIM method in accurately imputing a marker hinges on both input and output markers. When a latent biological relationship exists between the set of input markers and the output marker, our model excels in generating highly reliable imputed images for the output marker. It is worth noting that assessing imputed images on a pixel-by-pixel basis poses a challenge for markers exhibiting notably sparse responses, such as CD4, CD8, and PDL1. In such cases, both SSIM and MAE scores converge across MAXIM, BL-Zero, and BL-Mean methods. We applied MAXIM to multiplex immunofluorescence images featuring seven markers. Nevertheless, the versatility of our proposed method allows for seamless retraining and testing on images with varying marker counts. Likewise, it can be readily adapted for use with multiplex images obtained through diverse multiplex imaging technologies, including CODEX (2), MIBI (4), and others.

The MAXIM’s practical utility is threefold. First, the laboratories utilizing multiplexed images can seamlessly train an in-house MAXIM model using images devoid of staining issues. The trained model can then be employed to accurately impute markers in multiplexed images that are marred by staining problems. Second, MAXIM can serve as a valuable tool for quality control in newly generated multiplex images, aiding in the detection of staining failures. The strong correlation between imputed and real markers in new images will be an indicator of staining integrity. Finally, the interpretability of MAXIM provides the opportunity to uncover previously unknown latent biological relationships between different protein markers, leading to new insights in the field.

## Material and Methods

### Dataset

This study utilized a diverse dataset comprising tissue samples from four cancer types: Urothelial, Anal, Cervical, and Head and Neck. The dataset comprised 83 tissue samples from 27 cases derived from three separate studies, one specifically focused on Head and Neck cases (**Supplementary Table 1**). To acquire multiplex immunofluorescence (MxIF) image, formalin-fixed paraffin-embedded 5 *µm* sections from tissue samples were immune-stained using Opal 6plex kits, according to the manufacturer’s protocol (Akoya Biosciences), for a panel of DAPI, CD4, CD8, FOXP3, PDL1, KI67, and CK. Deparaffinizing, rehydration, epitope retrieval, and staining of slides were performed using Leica BOND RX Autostainer (Leica). The optimum staining condition for each antibody was determined using immunohistochemistry and single-immunofluorescence before combination. Details on antibodies, protocol, and opals used in this panel are described in **Supplementary Table 2**. The MxIF images were scanned at a high resolution of 40*×*, with a microns-per-pixel (mpp) value of 0.25. Among the 83 tissue samples, 29 samples from 5 cases were scanned entirely to produce whole slide MxIF images, while the remaining samples underwent separate scanning as regions of interest (ROI), resulting in ROI MxIF images of size 1396*×*1860 pixels. The images were unmixed using InForm version 2.5 software (Akoya Biosciences), enabling the identification and separation of weakly expressing and overlapping signals from the background autofluorescence.

The MxIF imaging dataset was divided into four sets for model training and evaluation: Train/Valid, Test-WSIs, Test-ROIs, and Test-HNSCC. The Train/Valid set consisted of 25 whole slide MxIF images selected from four cases (1 Urothelial, 1 Anal, and 2 Cervical), encompassing 14,476 images with dimensions of 1396*×*1860 pixels. To evaluate the model’s performance on unseen data, the Test-WSIs set comprised 1,920 images extracted from four whole slide MxIF images of a Head and Neck case. The Test-ROIs set was also created to assess the model’s robustness across diverse cases and tissue samples, including 1,097 images extracted from 9 cases and 28 tissue samples. Finally, a separate Test-HNSCC set was specifically prepared for evaluating the model’s performance on Head and Neck Squamous Cell Carcinoma (HNSCC) cases, consisting of 13 cases, 26 tissue samples, and a total of 623 images extracted solely from the HNSCC-focused study. These distinct data splits allowed for a comprehensive evaluation of the model’s generalizability and performance across different cancer types, tissue samples, and specific case studies.

### MAXIM model architecture

The MAXIM model is based on the U-Net architecture (26), which is an encoder-decoder network with skip connections. Let’s denote the input MxIF image as *X ∈* R*H×W ×N*, where *H* and *W* represent the height and width of the image, respectively, and *N* represents the number of markers. Each element *X*(*i, j, n*) corresponds to the intensity value of marker *n* at the pixel location (*i, j*). The MAXIM model aims to generate an imputed image *Y ∈* R^*H×W*^ for the output marker. The model consists of an encoder path and a decoder path connected by skip connections. The encoder path can be denoted as a function 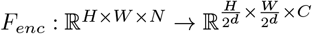 where *d* represents the depth of the encoder, and *C* represents the number of channels in the encoded feature map. Similarly, the decoder path can be represented as a function 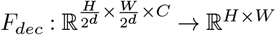, which impute the output marker image. Hence, the overall MAXIM model can be represented as a composite function *F* : ℝ^*H×W ×N*^*→*ℝ^*H×W*^, where *F* (*X*) = *F*_*dec*_(*F*_*enc*_(*X*)).

The MAXIM model was optimized using reconstruction loss, which involves minimizing the discrepancy between the imputed image and the ground truth image. In this case, *L*1 loss (mean absolute error) and *L*2 loss (mean squared error) were used as the reconstruction loss functions. Let’s denote the predicted imputed and the ground truth image as *Y ∈* R^*H×W*^ and *Y*_*gt*_ *∈* R^*H×W*^, respectively. The *L*1 loss is calculated as the mean absolute difference between the predicted and ground truth images:

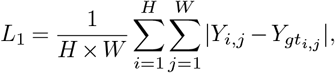

where *H × W* represents the total number of pixels in the image. Similarly, the *L*2 loss is computed as the mean squared difference between the predicted and ground truth images:

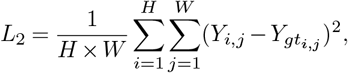

The optimization process of the marker imputation model involves minimizing the combined loss _*C*_, which is a linear combination of the L1 and L2 losses:

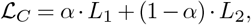

where *α* is weight parameter that control the relative importance of the L1 and L2 losses.

### MAXIM model with adversarial loss

The MAXIM model is also trained using an adversarial loss, following the training approach commonly used in conditional generative adversarial networks (27). The generator network, *G*, responsible for imputing the output marker image, adopts the same network architecture as described above, whereas the discriminator network *D* employed in the model is the same as the discriminator network utilized in the Pix2Pix network (28). The discriminator network input is a concatenated image of the multiplex input image with *N* markers *X ∈* R^*H×W ×N*^, along with either the ground truth output marker *Y*_gt_ or the generated output marker *G*(*X*). The discriminator network produces a 32 times smaller binary image as output. The discriminator network consists of six convolutional layers with a kernel size of 4 and a stride size of 2. Each convolutional layer is followed by a batch normalization operation and a leaky ReLU activation function, except for the first and last layers. The first layer does not include a batch normalization operation, and the last layer neither has a batch normalization operation nor a leaky ReLU activation. The discriminator’s objective is to discriminate and classify the input images into two categories: the first represents the multiplex input image with the ground truth marker, and the second represents the multiplex input image with the imputed marker. To achieve this, the discriminator predicts an output of all ones (close to one) for the first type of input image, indicating its high confidence that the image contains real values. Conversely, it predicts an output of zeros (close to zero) for the second type of input image, expressing its certainty that it contains imputed values. The discriminator model was optimized using the average cross-entropy loss (*ℒ*_D_) of images from both categories:

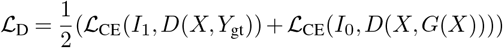

Here, *I*_1_ and *I*_0_ represent the images with pixel values as ones and zeros, respectively. The term *ℒ*_CE_(*I*_1_, *D*(*X, Y*_gt_)) computes the cross-entropy loss between the discriminator’s output when provided with the input image *X* and ground truth marker *Y*_gt_, aiming to maximize the probability of the discriminator correctly classifying them as real. Similarly, the term *ℒ*_CE_(*I*_0_, *D*(*X, G*(*X*))) calculates the cross-entropy loss between the discriminator’s output when provided with theinput image *X* and the imputed marker image *G*(*X*), aiming to maximize the probability of the discriminator correctly classifying them as imputed.

The generator network, *G*, is trained to impute marker values that closely resemble the ground truth markers and are challenging for the discriminator to differentiate. The generator network is optimized using the sum of weighted *L*1 loss (*L*_1_) and a cross-entropy loss. The generator loss is defined as:

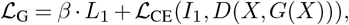

where *β* is the weight factor. The cross-entropy loss (*ℒ*_CE_) encourages the generator to generate marker predictions that the discriminator classifies as real. The overall loss ensures that the generator network is trained to produce accurate imputed marker values while also satisfying the adversarial objective.

### Training details

A separate model was trained for each marker using the remaining six as input. The images in the Train/Valid set were too large to be directly used for model training. Therefore, all images were divided into 24 equally sized patches of size 349*×*310. Then each model was trained using randomly cropped, flipped, and rotated regions of size 256*×*256 extracted from image patches. All models are optimized using Adam optimizer with a learning rate of 0.002. The values of *α* and *β* hyperparameters are empirically set to 0.5 and 100, respectively. The batch size was set to 64 for each training iteration. The maximum number of epochs was set to 200, while the minimum number of epochs was set to 50. A stopping criterion was implemented to prevent overfitting and ensure a stable training process. Training for each marker was stopped if the validation loss did not decrease for 25 consecutive epochs, indicating a potential plateau in model performance.

### Evaluation

The marker imputation models were evaluated using various evaluation metrics at the image level, considering the 1396*×*1860 pixel images. To assess the similarity between real and imputed marker images, the Structural Similarity Index (SSIM) was employed as a measure of structural resemblance. Additionally, the Mean Absolute Error (MAE) was calculated to quantify the pixel-level intensity differences between the real and imputed images. The evaluation of the marker imputation model considered the sparsity of marker images, which can vary depending on the marker type (nuclear, cytoplasmic, or membrane). Sparse images tend to yield higher SSIM scores and lower MAE values. The worst achievable SSIM score is 0, indicating no structural resemblance, while the worst MAE score is 255, denoting maximum average pixel-level intensity differences. To establish baselines for comparison, two sets of results were presented: one using images with all zeros and the other using images representing mean images of the input markers. These baselines provided the lowest achievable SSIM and highest MAE scores in the test sets. The MAXIM’s performance is expected to surpass these baseline results, demonstrating its ability to impute more accurate and realistic marker images. In addition to evaluating the MAXIM’s performance based on pixels, we also conducted a cell-level assessment to mitigate potential biases arising from numerous zero or nearzero pixel values in both real and imputed marker images. HALO image analysis software (Indica Lab), specifically the automated Highplex FL module, was utilized to calculate the mean expression of each cell in the real and imputed markers. Initially, the software identified and segmented the cell nucleus using the real DAPI channel and then expanded the segmented region using a heuristic approach to encompass the entire cell, including the cell membrane. Subsequently, it determined the average intensity or expression within each cell for both the real and imputed marker images.

To quantify the intensity differences between the real and imputed marker cells, we computed the MAE at the cell level by comparing the mean cell expression values. Additionally, we assessed the model’s performance in cell classification by utilizing the mean cell expression of the real marker. The HALO software facilitated this classification task, wherein an expert-defined threshold was set. If the mean expression of the pixels belonging to a cell exceeded the threshold, the cell was classified as positive for that marker. Using the resulting cell labels and the mean cell expression of the imputed marker images, we generated precision-recall curves to illustrate the trade-off between precision and recall. The average precision (AP) score served as a summary measure of the model’s ability to distinguish between positive and negative cells based on the imputed marker values.

### Model Interpretability

Model interpretability was explored by calculating aggregated gradient-based image attributions (25) of each marker in the input images to the imputed marker image. These attributions, derived from the model’s internal computations, could take both positive and negative values, indicating the contribution of each pixel to the imputed marker image. The absolute sum of attributions was computed for each marker to assess the overall attribution of each input marker for a given output marker. This yielded a measure of the importance or influence of each marker in the model’s decisionmaking process. Furthermore, the aggregated attributions of each marker were converted into percentage values by dividing the total absolute attribution of the input image. This conversion provided a more interpretable representation of the relative significance of each marker in the context of the entire image. In addition to calculating aggregated gradientbased pixel attributions, the correlation between pairs of input markers’ attributions was also computed to identify any patterns of co-occurrence in marker imputation. This analysis aimed to examine whether certain input markers exhibited similar attribution patterns or if there were any interdependencies between the attributions of different markers.

To validate the attribution of individual input markers to the imputed marker image, we conducted experiments using two sets of new MAXIM models. Each set consisted of a subset of markers with either high or low attribution scores. In one set, the input images were modified to exclude the markers with high attribution scores. Conversely, the input images were modified in the other set to exclude the markers with low attribution scores. The intention behind this approach is that models trained without markers with low attribution scores should ideally perform consistently better compared to the other set of models. We calculated the MAE metrics for each set and compared them with the results obtained from the original marker imputations. This analysis allowed us to evaluate the significance of the attributions of input markers with high and low scores and determine their influence on the model’s performance.

### Ablation study

In order to investigate the influence of training set size on the performance of the MAXIM model, we conducted an ablation study by training four new MAXIM models using varying proportions of the available training images. Specifically, we randomly selected 75%, 50%, 25%, and 10% of the training images for each model. By training models with reduced training data, we aimed to understand how the availability of labeled data affects the model’s ability to accurately impute markers in MxIF images.

### Computational hardware and software

We train all models on a system with Intel Core i9-10920X CPU (central processing unit) and an NVIDIA GeForce RTX 3090 GPU (graphics processing unit). All models were implemented in Python (3.9.13) using PyTorch (1.13.1) and TorchVision (0.14.1) as the primary deep-learning packages for the development and training of deep learning models. Numerical computations and data manipulation were carried out using Numpy (1.12.5) and Pandas (1.4.4), respectively. The Scipy (1.9.1) was utilized to calculate evaluation metrics such as MAE, precision-recall curve, and average precision, while Scikit Image (0.19.2) was used to calculate the SSIM between two images. Model interpretability was achieved through the application of Captum (0.6.0). The data visualization aspect of the research employed Matplotlib (3.5.2) and Seaborn (0.11.2), enabling the creation of informative visual representations. Setuptools (63.4.1) assisted in managing library dependencies for pip package creation.

## Data availability

Data will be shared upon reasonable request. Any request for data by qualified scientific and medical researchers for legitimate research purposes will be subject to NCI and the NIH Policy for Data Management and Sharing. All requests should be submitted in writing to NCI.

### Code availability

All code and scripts for reproducing the experiments are accessible at https://github.com/mahmoodlab/MAXIM. Detailed instructions are provided in the README for easy setup and replication of results.

## ACKNOWLEDGEMENTS

This project has been funded in whole or in part with Federal funds from the National Cancer Institute, National Institutes of Health, under Contract No. 75N91019D00024. The content of this publication does not necessarily reflect the views or policies of the Department of Health and Human Services, nor does mention of trade names, commercial products, or organizations imply endorsement by the U.S. Government. Support was part of the Frederick National Laboratory’s Laboratory Directed Exploratory Research (LDER) Program.

## AUTHOR CONTRIBUTIONS

F.M., G.Z., Y.L, and M.S. conceived the study and designed the experiments. M.S. performed the experimental analysis. W.L., S.B., C.A., J.S., and G.Z curated the dataset. K.C. and W.L. process the data in HALO. M.S., W.L., G.Z., H.A.S., J.L.G, and F.M. analyzed the results. M.S., S.R., and F.M. prepared the manuscript with input from all co-authors. G.Z., F.M., and H.A.S. supervised the research.

### CONFLICT OF INTERESTS

The authors declare no competing interests.

---

**Extended Data Figure 1.**
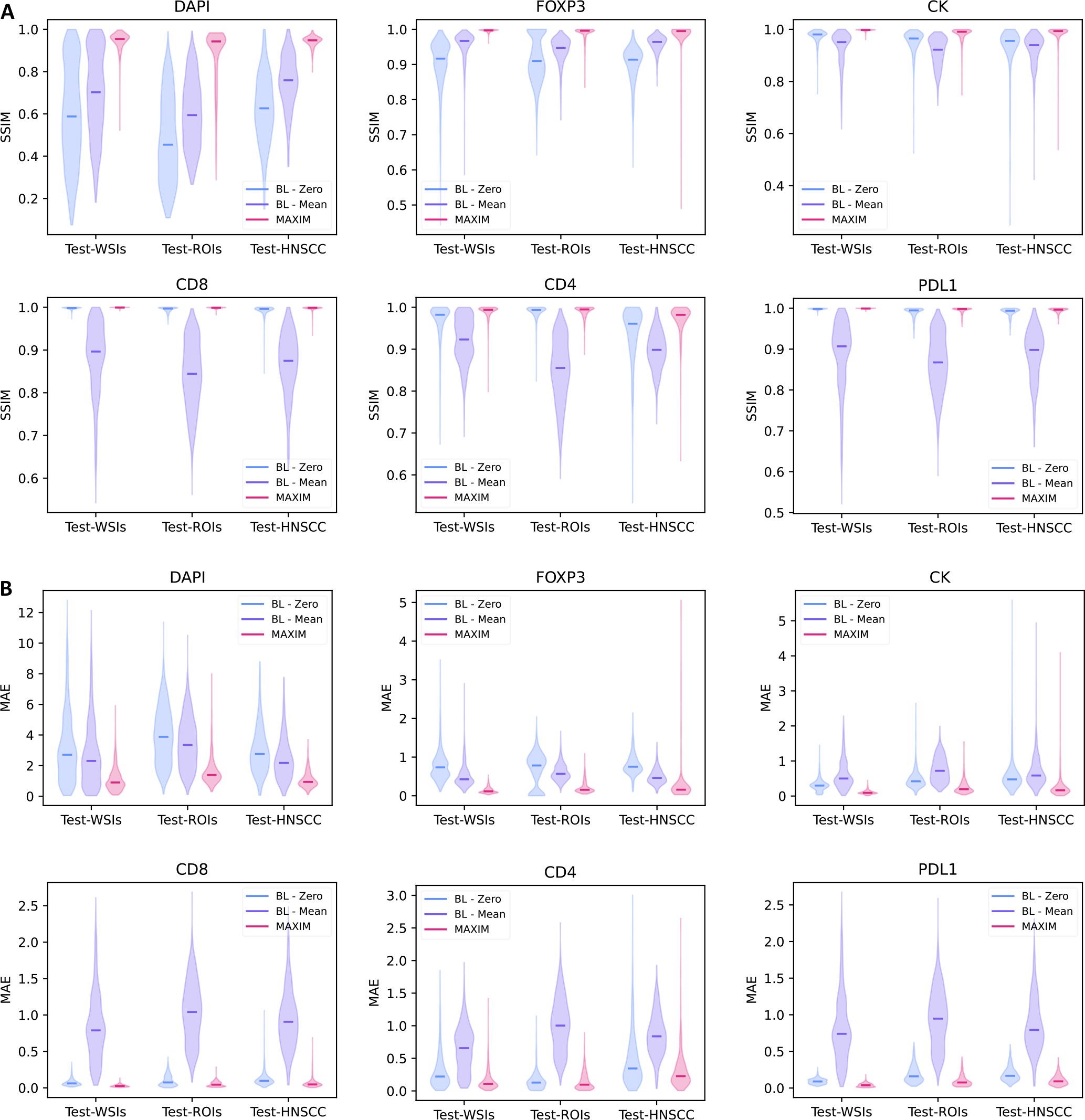
The quantitative assessment of MAXIM’s marker imputation ability for six different markers. **(A) & (B)** present the results using structural similarity index (SSIM) and mean absolute error (MAE) metrics across three test sets, respectively. The width of each violin represents the density of data points (images) with corresponding SSIM/MAE scores, and the solid line within each violin indicates the median SSIM/MAE score. BL-Zero and BL-Mean are pseudo models that provide baseline MEA and SSIM scores in multiplex images with high sparsity (CD8, CD4, CK, and PDL1) and structural similarity (DAPI and FOXP3) across markers. BL-Zero generates an imputed image with zero values, while BL-Mean creates an imputed image using the mean values of the input markers.

**Extended Data Figure 2.**
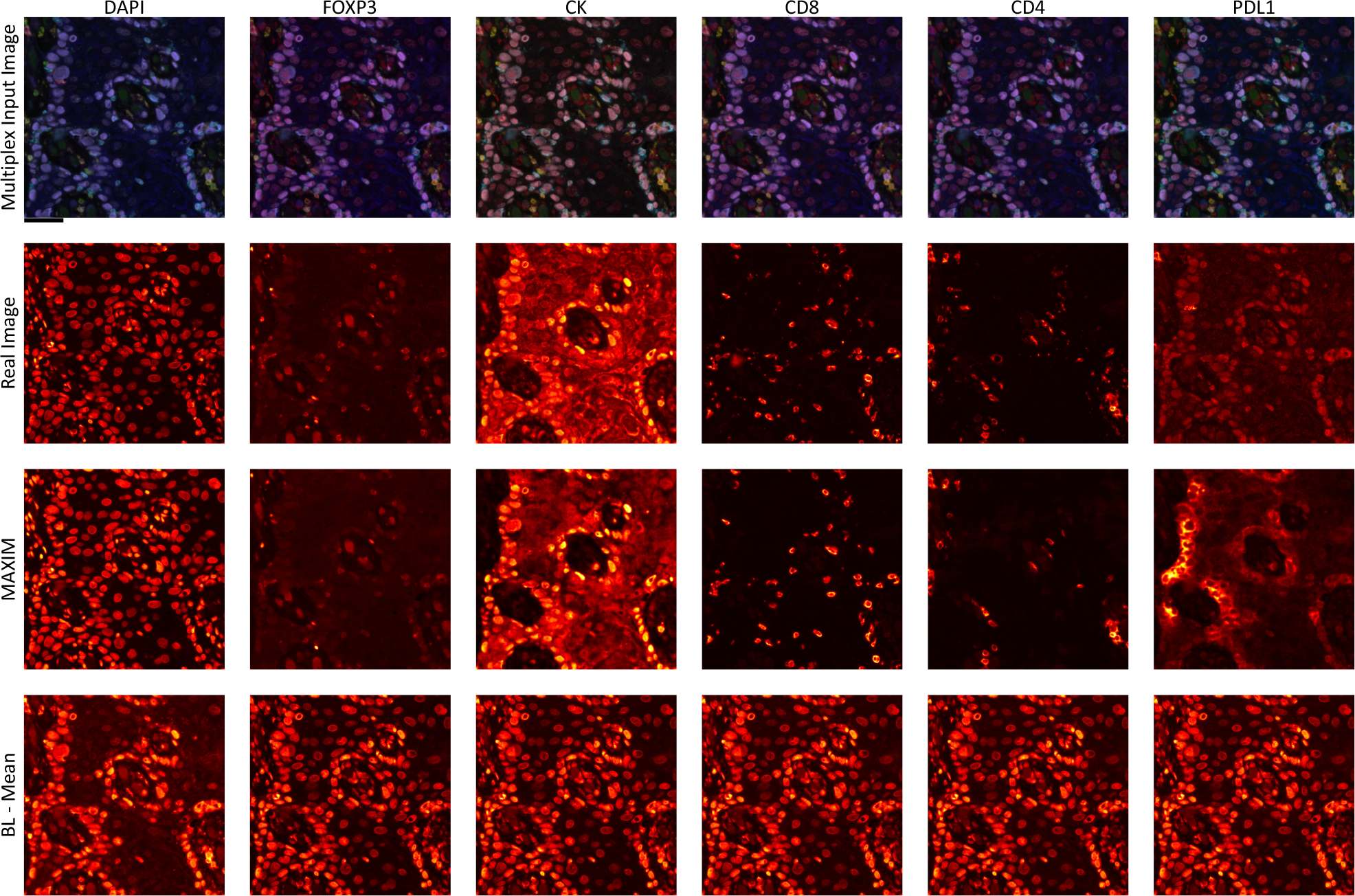
Visual results of six different marker imputation models. The results are shown for the same multiplex image with seven markers: DAPI (red), FOXP3 (green), CD4 (yellow), CD8 (cyan), PDL1 (magenta), KI67 (White), and CK (blue). Scale bar: 50 *µm*. The first row shows the input images consisting of six markers, excluding the target marker (column name). The second row shows the real images of the target markers. The third and fourth rows show imputed images using MAXIM and BL-Mean models.

**Extended Data Figure 3.**
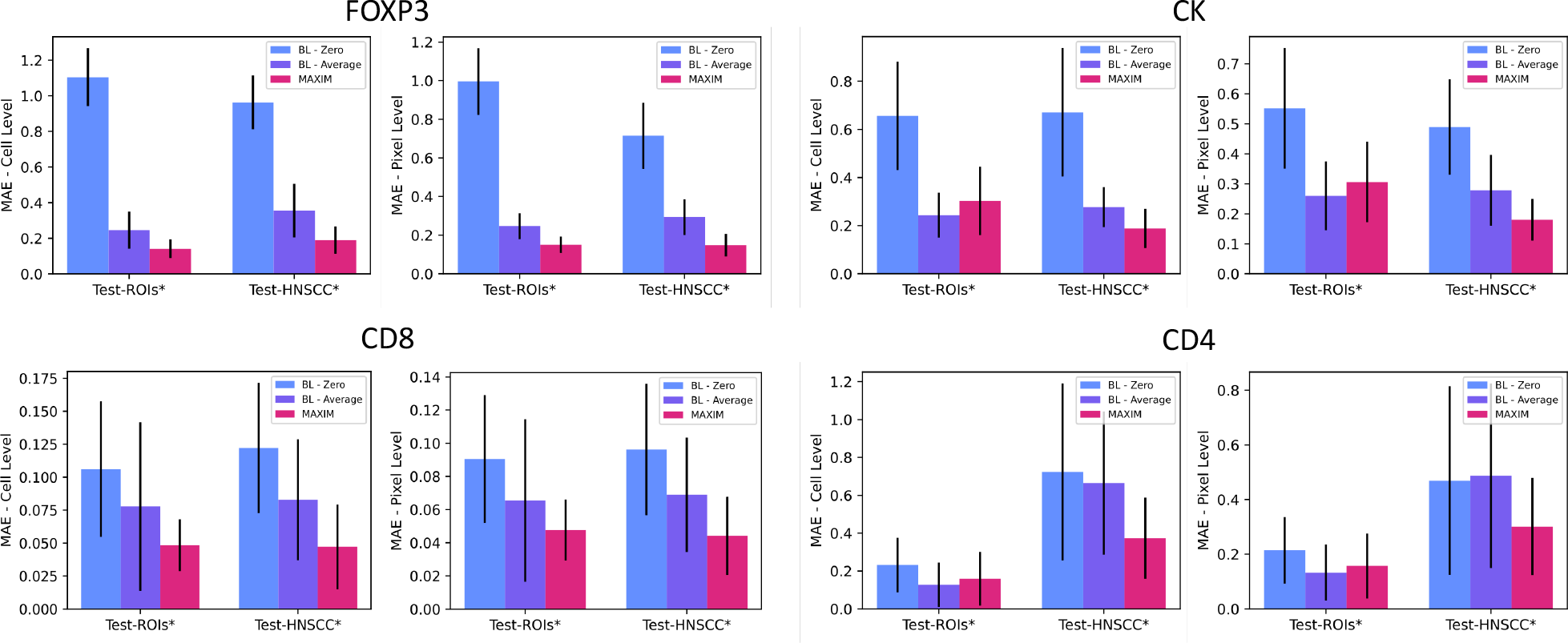
Evaluation of MAXIM at cell and pixel level on subsets of Test-ROIs and Test-HNSCC sets for six different markers. The difference in the mean expression of each cell is used for cell level evaluation. The height of the bars indicates the average absolute error, whereas error bars represent the standard deviation.

**Extended Data Figure 4.**
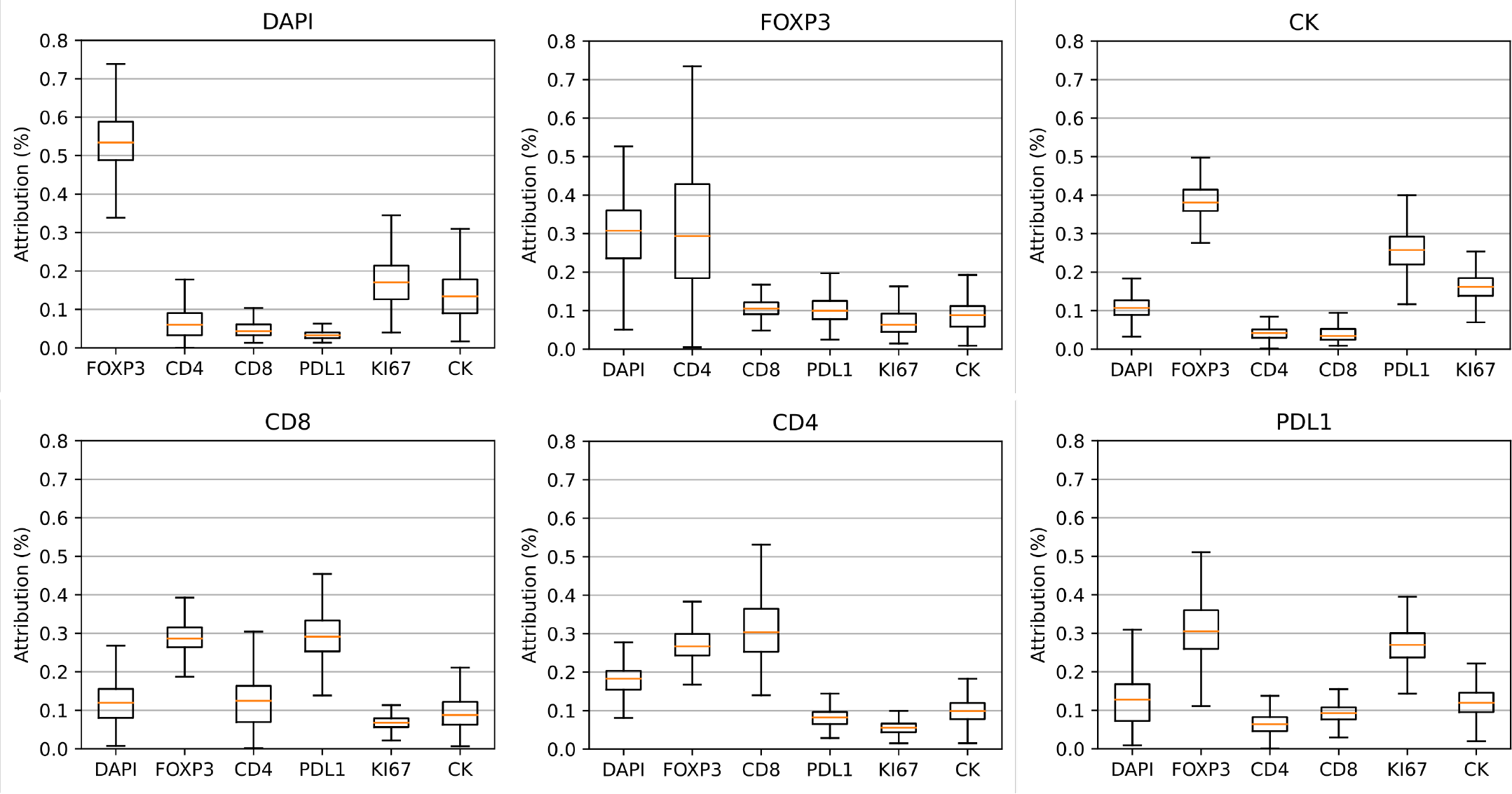
Model interpretability analysis using aggregated gradient method. The boxplots show the distribution of gradient-based attribution of each input marker towards output marker imputations. The line inside the box represents the median, the edges of the box represent the interquartile range, and the whiskers extend to 1.5 times the interquartile range to represent the range of the attribution values.

**Extended Data Figure 5.**
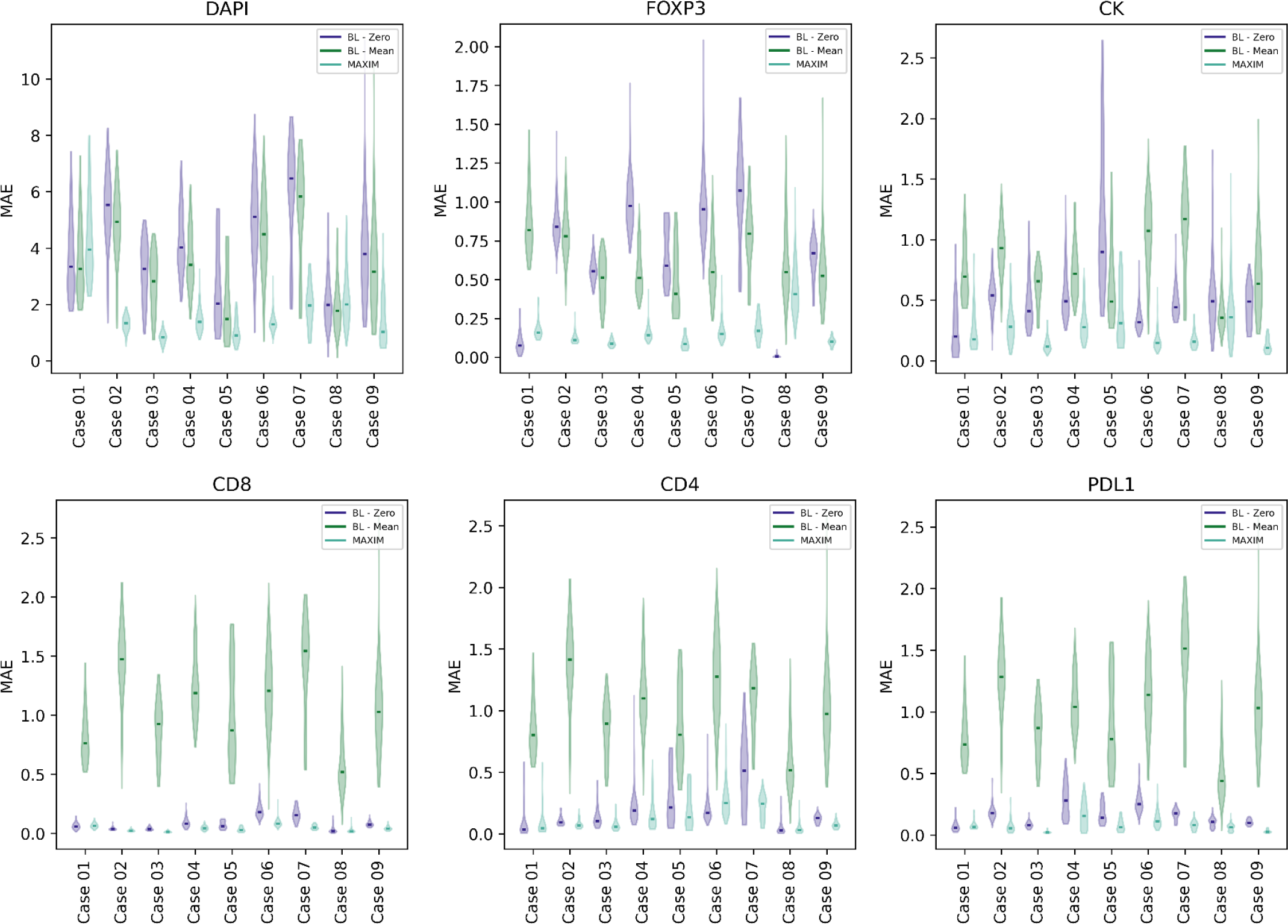
Evaluation of MAXIM model at case level on Test-ROIs set. The width of each violin represents the density of data points (images) with corresponding mean absolute error (MAE) scores, and the solid line within each violin indicates the median MAE score.

**Extended Data Figure 6.**
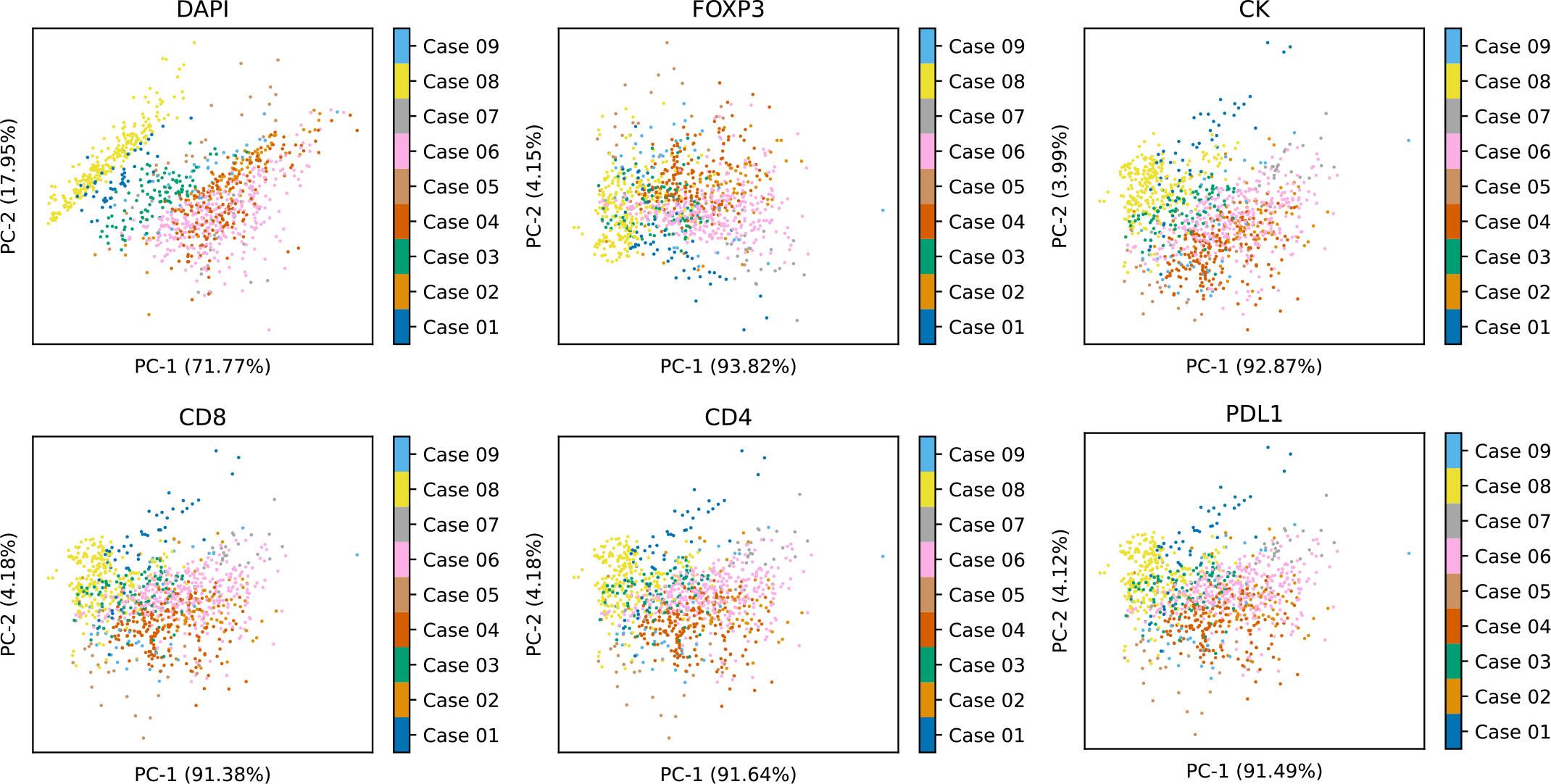
Two-dimensional projection of input images to visualize the case level differences in data distribution for different MAXIM models. Each plot showcases the projections achieved using principal component analysis, which are based on the mean intensities of each marker in the input images. Each dot within the subplots represents an individual image from the Test-ROIs set, and the color of the dots indicates the corresponding case to which the image belongs.

## Notes

https://github.com/mahmoodlab/MAXIM

### Competing Interest Statement

The authors have declared no competing interest.

